# The molecular mechanism of temperature-dependent phase separation of Heat shock factor 1

**DOI:** 10.1101/2024.08.20.608825

**Authors:** Qiunan Ren, Linge Li, Juan Li, Chaowei Shi, Yujie Sun, Xuebiao Yao, Zhonghuai Hou, ShengQi Xiang

## Abstract

Heat shock factor 1 (HSF1) is the critical orchestrator for a cell responding to heat shock, and its dysfunction is linked to cancers and neurodegenerative diseases. HSF1 undergoes phase separation upon heat shock, and its activity is regulated by various post-translational modifications (PTMs). Despite its importance, the molecular details underlying HSF1 phase separation, temperature sensing, and post-translational modifications (PTMs) regulation remain poorly understood. Here, we discovered that HSF1 exhibits temperature-dependent phase separation with a lower critical solution temperature (LCST) behavior due to entropy contribution from solvent molecules, providing a new conceptual mechanism accounting for HSF1 activation. We employed a synergistic approach combining coarse-grain simulation and nuclear magnetic resonance spectroscopy to reveal the residue-level molecular details of the interactions driving the phase separation of wild-type HSF1 and its distinct PTM patterns at various temperatures. The identified interaction sites were further validated with biochemistry assays and mapped interface accounts for HSF1 functions reported. Importantly, the amino acid substitution experiment reveals the molecular grammar for temperature-dependent HSF1 phase separation is species-specific and physiologically relevant. These findings delineate chemical code that integrates protein PTM patterns with accurate phase separation for body physiological temperature control in animals.

## Introduction

The heat shock response (HSR) is a ubiquitous cellular response that plays a vital role in dealing with various stress conditions to maintain protein homeostasis. Heat shock factor 1 (HSF1) serves as the central regulator in HSR, orchestrating numerous physiological processes^1^. At the physiological temperature, HSF1 is bound with molecular chaperones, keeping it monomeric and inactive^2,3^. However, when cells are exposed to high-temperature stress, the monomer HSF1 aggregates and translocates to the nucleus, where it promotes the transcription of various regulatory factors that help maintain the homeostasis of the intracellular environment^4^.

The fine-tuning of HSF1 activation is critical, as any malfunction of regulation will result in disease. Elevated HSF1 activity has been found to inhibit apoptotic genes and activation of cell metabolism genes, contributing to the proliferation of cancer cells^5–7^. Conversely, in Huntington’s disease, reduced HSF1 decreases the expression of molecular chaperones, leading to neuronal dysfunction^8,9^. Previous studies have identified various post-translational modification (PTM) sites within HSF1 that modulate its function in both directions *in vivo*^10^ (Fig 1a). These PTMs have been classified and summarized according to their effects on HSF1 function, and two representative mutants, M1 and M2, have been proposed to mimic the PTM patterns of activating and inhibiting protein functions, respectively^10^.

**Figure 1.**
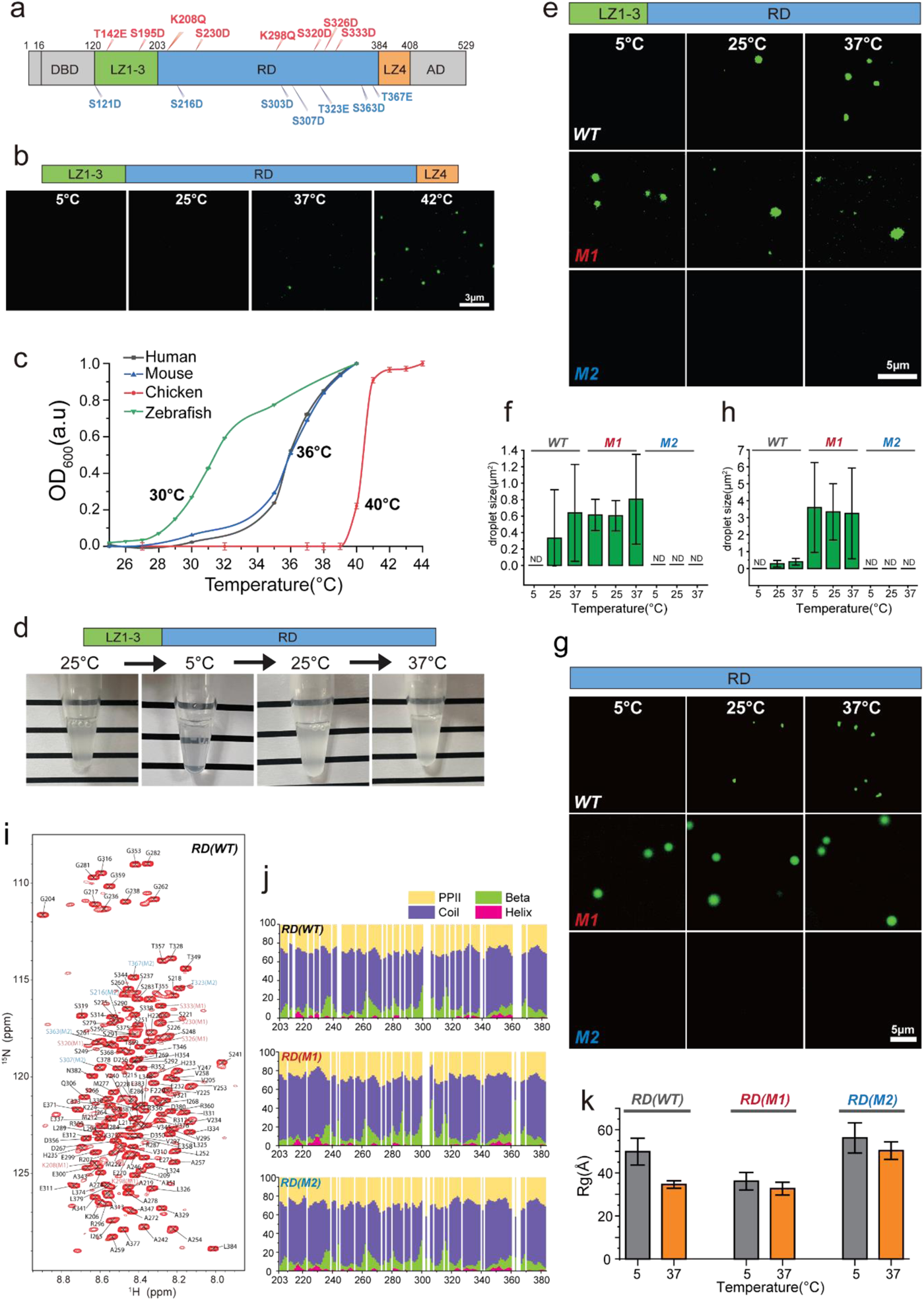
The RD region drives temperature-dependent phase separation. **a** Schematic representation of human HSF1 domain organization. Mutants M1 (red) and M2 (blue) were designed to mimic two distinct PTM patterns in HSF1, with S to D and T to E mutations used to mimic constitutive phosphorylation. **b.** Fluorescence images of droplets formed by 1 µM LZ1-3-RD-LZ4 at different temperatures (5°C, 25°C, 37°C, and 42°C ). Scale bars, 3 µm. **c.** The absorption values at 600 nm of WT LZ1-3-RD-LZ4 varied with temperature. The turbidity curve showed the protein underwent phase separation within a narrow temperature range. **d.** Photographs of WT LZ1-3-RD showed its reversible phase separation, which was promoted at 25°C and 37°C and vanished at 5°C . **e.** Representative images of fluorescence droplets formed by 1 µM WT LZ1-3-RD and mutants (M1 and M2) at 5°C , 25°C , and 37°C . Scale bars, 5 µm. **f.** Droplet sizes from the different treatment groups in **e**. ND, not determined. **g.** Representative fluorescence images of droplets formed by 120 µM WT RD domain and mutants (M1 and M2) at 5°C , 25°C , and 37°C . Scale bars, 5 µm. **h.** Droplet sizes from the different treatment groups in **g**. ND, not determined. **i.** The ^1^H-^15^N HSQC spectrum and backbone assignments of the WT RD domain. The mutation sites are in red for M1 and blue for M2. **j.** Secondary structure populations of WT RD, M1 and M2 obtained from the delta2D method using experimentally assigned backbone chemical shifts. **k.** Rg values of WT RD, M1, and M2 mutants were measured by SAXS at 5 ℃ and 37 ℃.

A model of HSF1 activation based on protein oligomerization has been proposed, in which HSF1 is activated by forming a trimer and inactivated in its monomeric form^11^. In this model, the intramolecular interactions between two leucine zipper regions, LZ1-3 and LZ4, work as an autoinhibition mechanism to keep HSF1 as a monomer. When the temperature is elevated, LZ1-3 disassociates from LZ4 and mediates intermolecular oligomerization^11^. However, this model is not in line with the observation that most of the PTM sites that modulate HSF1 activity are not primarily located in the leucine zipper regions but instead concentrated in the disordered regulatory domain (RD)^10,12,13^.

Phase separation of biomolecules has been extensively observed throughout the cell life cycle and is closely related to many physiological functions. Particularly in the cell nucleus, many phase separation phenomena in the transcription process have been reported^14–16^. Recently, the phase separation of HSF1 has been reported^17,18^. HSF1 droplets have been observed in tissues of various cancers, and these droplets are fluid^19^. Phase separation of HSF1 stimulates the upregulation of several HSP genes in cells, indicating that phase separation of HSF1 plays a vital role in HSR^13^. Subsequently, researchers have found that PTMs significantly impact the phase separation ability of HSF1^13^. The two representative mutants mimicking distinct PTMs patterns, M1 and M2, which activate or suppress HSF1 activity in cells, also up- or down-regulate the phase separation properties of HSF1 *in vitro* and in cells^13^.

The concept of phase separation represents a fresh avenue for understanding the temperature sensing and activation of HSF1. Since the HSF1 activates upon the heat shock and its function is related to its phase separation, exploring the relationship between temperature and HSF1 phase separation is worthwhile. Furthermore, the underlying molecular mechanisms, particularly the high-resolution details involved in the temperature and PTM regulation of phase separation, remain to be elucidated, which requires accurate characterization of the atomic level interactions.

To gain insights into the molecular mechanisms underlying phase separation, various computational approaches alongside NMR spectroscopy have been employed in previous research efforts^20–22^. While all-atom molecular dynamics (MD) simulations have successfully characterized protein conformations and interactions^23,24^, a wide range of coarse-grained (CG) models have been developed to understand the complete phase behavior and reproduce the condensation process^20,25–27^. Among these models of diverse resolutions, residue-based CG models have proven to be particularly useful in explaining the sequence-dependent phase separation of intrinsically disordered regions (IDRs), which retain physio-chemical features of the protein sequence and affordable simulation efficiency^28,29^. However, although various CG force fields have been developed^30,31^, such models have proven difficult to generalize to predict the phase behavior of diverse sequences^28^. Therefore, incorporating atomic-level information obtained from high-resolution computation or experimental data may be indispensable to improve the accuracy of the CG model^32,33^. NMR spectroscopy has been shown to be a valuable tool for studying IDRs, providing high-resolution information on conformation, dynamics, and interactions of biomolecules before and after phase separation^20–22,34–38^.

In this work, we explored the temperature-dependent properties of the HSF1 phase separation, which exhibited an LCST-type behavior attributed to its IDR domain and matched with animal body temperature. With an integrated approach of NMR spectroscopy and simulation, the molecular mechanisms of phase separation and the differential regulation of the distinct PTM patterns were revealed with residue resolution. The identified interacting regions are shown to determine the phase separation temperature and fit with previous cell biology results.

## Results

### The phase separation of HSF1 is temperature-dependent and driven by the intrinsically disordered RD domain

It has been previously reported that the phase separation of HSF1 is promoted *in vivo* at elevated temperatures, and the phase separation ability was attributed to the region containing only the leucine zippers (LZ1-4) and RD domains^13^(Fig. 1a). Here, we investigated the phase separation behaviors of various purified HSF1 fragments *in vitro,* ranging from 5 ℃ to 42 ℃. At a protein concentration of 1 µM, the LZ1-3-RD-LZ4 fragment began to form droplets at 37 ℃, with more pronounced phase separation observed at 42 ℃ (Fig. 1b). The turbidity-temperature curve of the wild-type LZ1-3-RD-L4 indicated that phase separation occurs over a very narrow temperature range (Fig. 1c). HSF1’s phase separation is highly sensitive to temperature changes, making HSF1 an ideal temperature sensor.

Since the body temperatures vary among animal species and the HSF1 is activated above the physiological temperature, the phase-transition temperatures of HSF1 were speculated to exhibit differences across species. To test this hypothesis, we compared the temperature-dependent phase transition behaviors of HSF1 proteins from several animal species with distinct body temperatures, including human, mouse, chicken, and zebrafish (*Danio rerio*) (Fig. 1c). Our results showed that the HSF1 proteins from humans and mice started to show phase separation around 37 ℃, corresponding to the body temperature of mammals^39^. In contrast, the chicken HSF1 only underwent phase separation when the temperature was above 40℃, consistent with the higher body temperature of chickens (∼41℃)^40^. The optimum temperature for zebrafish is 28℃^41^, while the HSF1 of zebrafish showed a phase transition in the range between 28 and 32℃. Therefore, our findings suggest that the phase transition temperature of HSF1 is in good correlation with animal body temperatures (Fig. S1a).

When the LZ4 fragment was not included, the phase separation of LZ1-3-RD occurred over a wide range of temperatures, with droplets already apparent at 25 ℃ and further pronounced at higher temperatures (Fig. 1d and 1e). The phase separation of LZ1-3-RD supported the negative regulatory role of LZ4, as previously reported^13^. Notably, the phase separation was also reversible when the temperature was lowered again to 5 ℃ from 25 ℃ (Fig. 1d). Moreover, some droplets showed irregular shapes at 37 ℃, suggesting a liquid-to-gel transition.

Then, the two PTM mimicking mutants employed in the previous studies, M1 and M2, were also subjected to phase transition analysis. Consistent with the previous report, the mutant M1 mentioned above was more prone to phase separation than the wild type, while the mutant M2 lost its phase separation ability in the experimental setting at all temperatures tested. Interestingly, the phase separation behavior of both M1 and M2 did not change with temperature, contradicting the wild-type protein. (Fig 1e and 1f).

To further define the core region responsible for the phase separation of HSF1, we generated LZ1-3 and RD constructs separately (Fig. 1a) and assessed their respective phase separation ability. The LZ1-3 region was reported to be prone to oligomerization ^11^. Cross-linking and size-exclusion chromatography coupled with multiple angle light scattering (SEC-MALS) revealed that LZ1-3 was indeed in an oligomeric state in solution, consistent with previous results (Fig. S1b and S1c). However, even at a concentration of 500 µM, no droplets of LZ1-3 were observed. In contrast, the RD region was found to be sufficient to undergo phase separation by itself and retained the temperature dependency of the longer constructs, although the threshold concentrations were much higher than those of the longer LZ1-3-RD construct, 120 µM vs. 1 µM for LZ1-3-RD (Fig. 1g and 1h). These findings suggest that the RD region plays a crucial role in the phase separation of HSF1, matching the observation that most of the mutants in M1 and M2 were located within the RD domain rather than the helical LZ1-3 and LZ4 regions.

After establishing the RD domain as the core for HSF1 phase separation, we employed solution-state NMR to reveal the high-resolution structural information of the RD domain. First, the ^1^H-^15^N HSQC spectrum of the wild-type RD region was recorded, which clearly displayed the IDR properties of the narrow peak distribution (Fig. 1i). The backbone assignment was achieved with routine triple-resonance experiments covering 142 out of 153 non-proline residues. Most residues displayed secondary chemical shifts of less than 0.5 ppm, indicating a lack of stable secondary structure elements (Fig. S2a). The IDR nature of the RD domain was also confirmed by Delta2D analysis of the backbone chemical shifts (Fig. 1j). The hetNOE values of the backbone amide groups were sensitive to nanosecond range dynamics, with hetNOE values in the wild type being less than 0.5, demonstrating the flexible backbone. All these findings demonstrated that the RD region is an intrinsically disordered domain (Fig. S2b).

The backbone assignments of M1 and M2 were successfully achieved with all 153 non-proline residues in both cases (Fig. S3a and S3b). The secondary chemical shifts of M1 and M2 indicated that the mutations did not induce any local secondary structural propensity (Fig. 1j and S2a). The backbone hetNOE values for the two mutants also indicated that their backbones had the same degree of flexibility as the wild type (Fig. S2b). Furthermore, the mutations had no significant effects on the backbone T1 and T2 values (Fig. S2c and S2d).

The chemical shift temperature coefficients of every amide proton were calculated for both high (20 ℃ - 25 ℃) and low (5 ℃ - 10 ℃) temperature ranges. The ^1^H-^15^N HSQC spectra of WT RD and two mutants (M1 and M2) were obtained at various temperatures ranging from 5 ℃ to 25 ℃, and the backbone assignments were transferred from 5 ℃ to different temperatures. The temperature coefficients of amide protons are indicative of hydrogen bond formation. All values calculated in this study were below the threshold value of a stable intramolecular hydrogen bond, highlighting the characteristics of an IDR. Notably, the wild type exhibited a consistent and noticeable decrease as the temperature increased, while the two mutants showed no observable differences between these two temperature ranges (Fig. S4). The slight but discernable negative shifts implied the temperature-dependent conformation changes of the protein itself or the difference in the hydration shell.

Evidence suggests that the compactness of a single chain is well correlated with its phase separation ability^42,43^. Thus, we employed small-angle X-ray scattering (SAXS) to measure the dimensions of monomeric RD domains in a dilute solution at 5 ℃ and 37 ℃ to correlate the different phase separation behaviors with the molecular sizes. The radii of gyration (Rg) for three RD versions (wild-type, M1, and M2) were found to be different, with values of 49.9 Å, 36.1 Å, and 56.3 Å at 5 ℃ for wild-type, M1 and M2 mutants, respectively, in agreement with their phase separation ability. Furthermore, the molecule volume of the wild-type protein displayed a temperature-dependent change, with the Rg decreasing from 49.9Å to 34.6 Å from 5 ℃ to 37 ℃. In contrast, the Rg of M1 and M2 showed much more minor changes with temperature, with values of 32.7Å and 50.3Å, respectively, at 37 ℃ (Fig. 1k).

### HSF1 undergoes a phase separation with a lower critical solution temperature

In order to gain a comprehensive picture of the phase behavior of HSF1, we obtained binodals experimentally by measuring the concentrations of coexisting diluted and condensed phases between the temperature of 5 ℃ and 35 ℃ using both wild-type and M1 mutant LZ1-3-RD. In our experimental setting (200 µM protein in 25 mM Na-P, pH 7.0, 150 mM NaCl, and 2% PEG 4000), it was observed that the wild-type protein only demonstrated phase separation above ∼5℃. The binodal points obtained from the experiments were consistent with the observations shown in Fig. 1 and suggested that the phase separation propensity of WT LZ1-3-RD is weakened as the temperature decreases, with a critical temperature of approximately 1℃, indicating lower critical solution temperature (LCST) phase behavior (Fig. 2a). It is worth noting that most disordered proteins, whose driving force of phase separation is typically the enthalpy of multivalent interaction, usually exhibit upper critical solution temperature (UCST) behavior. LCST, on the other hand, is not commonly observed in interaction-induced biological condensation despite a few recent reports ^44–46^. Since the entropy of proteins usually decreases from mixed to demixed states, it is speculated that entropy gain can only come from solvent molecules. Protein surface at hydrophobic regions leads to restricted water molecules, which limits the degrees of freedom and decreases entropy (Fig. 2b left). When the protein undergoes phase separation, some associated water molecules are released from the protein due to the decrease in the surface, resulting in positive entropy gain (Fig 2b right).

**Figure 2.**
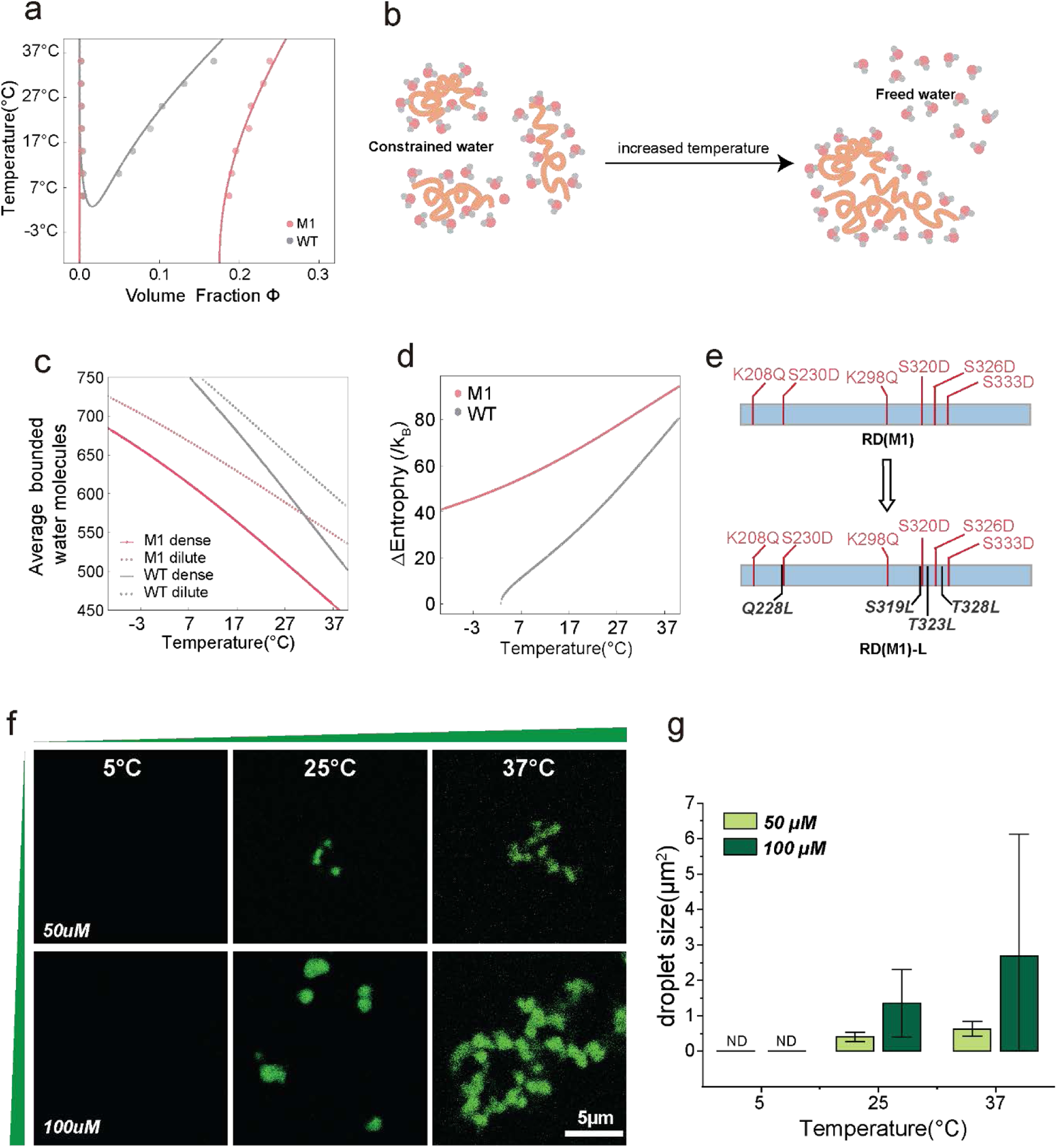
HSF1 undergoes an LCST-type phase transition attributed to the solvent entropy changes. **a.** The phase diagram of LZ1-3-RD for wild-type and M1. The lines are the fitting results of mean-field model fitting, while the dots are experimental results. **b.** Schematic illustration of the entropy-driven phase separation due to the change in associated water. The total water molecules are the same before and after phase separation. Please note that the scheme exaggerated the degree of water loss for clarity. **c.** The average restricted water molecules per chain decreased monotonically with temperature. **d.** Entropy difference upon phase separation with respect to temperature. The entropy gain was positive after phase separation, confirming the entropy-driven mechanism. The temperature sensitivity of the wild-type is higher than that of M1, which is attributed to the faster increase in entropy gain. **e.** Sequence scheme of RD(M1) and RD(M1)-L mutants. The mutations in the M1 for mimicking PTM were labeled in red, while the hydrophobicity compensation mutations are in black. **f.** Representative fluorescence images of 50 µM and 100 µM RD(M1)-L at 5°C, 25°C, and 37°C. Scale bars, 5 µm. **g.** Droplet sizes from the different treatment groups in **f**. ND, not determined.

In order to test our hypothesis regarding the entropy-driven model for HSF1, a mean-field model from polymer science is adapted ^47^, which considered the integration of entropy and enthalpy change caused by the formation of restricted water with a protein segment, in addition to Flory-Huggins’s terms (please refer to the Supplementary Material for details). The binodal points of wild-type LZ1-3-RD were well-fitted into a phase diagram with LCST using a simple mean-field model (gray line in Fig. 2a; see also Table S1). By reducing the entropy gain for restricted-water formation, M1 LZ1-3-RD data could also be fitted into a phase diagram with LCST using a simple mean-field model with solvation despite the critical temperature being far below the freezing temperature (red line in Fig. 2a, see also Table S1). Therefore, the LZ1-3-RD M1 mutant was observed to be always phase-separated within the experimental temperature range.

Since the positive entropy gain implies the release of restricted water molecules, we could examine the underlying phase separation mechanism by calculating the average restricted water per protein. Analysis using the fitted model revealed that average restricted water molecules per protein decrease monotonically with temperature, and the difference in this number between dense and dilute phases enlarges as the temperature rises (Fig. 2c). Interestingly, the restricted-water number of WT LZ1-3-RD declines faster than that of M1 LZ1-3-RD, indicating the higher sensitivity to temperature of WT LZ1-3-RD. Calculation of the entropy change yielded a positive entropy gain after phase separation, which confirmed the entropy-driven mechanism (Fig. 2d). Similarly, the entropy gain of WT LZ1-3-RD was observed to grow more rapidly than that of M1 LZ1-3-RD, which explains the higher temperature sensitivity in the phase separation of the wild-type. Thus, by fitting a mean-field model, we have identified the critical parameter related to solvent entropy responsible for the LCST phase behavior of HSF1 proteins. Our results suggest that the LCST phase behavior of WT is entropy-driven due to the presence of protein-bounded water molecules.

Since the phosphorylation (and the mutant mimics it) reduces the hydrophobicity^48–50^, the M1 is less hydrophobic than the WT, which results in less restricted water on its surface and a lack of LCST-type phase transition. To test this model, we compensated the hydrophobicity loss of phosphorylation by making additional mutations of more hydrophobicity, such as Ser to Leu, around the phosphorylation sites (Fig. 2e). The resulting protein, dubbed as RD(M1)-L, indeed showed an LCST behavior similar to WT (Fig. 2f and 2g). Hence, we managed to introduce the temperature dependency into the M1 mutant by manipulating the hydrophobicity of the protein.

### Intermolecular interactions mediate the phase separation of HSF1

To gain high-resolution insights into the phase separation mechanism, we conducted NMR titration experiments to probe the intermolecular interactions among the RD domains. Two samples were prepared: a reference sample containing only 0.1 mM ^15^N-labeled RD domain, and another sample containing 0.1mM ^15^N-labeled RD domain along with 1mM natural abundance RD domain (Fig. 3a). The NMR signals of both samples were from the same amount of ^15^N-labeled RD proteins, and the addition of a 10-fold excess RD protein led to changes in NMR results via intermolecular interactions. To exclude influences of viscosity by increased protein concentration, the diffusion coefficient (*D_T_*) of 1,4-Dioxane in 0.1mM and 1mM RD were measured using DOSY (Fig. S5a), which showed marginal differences (∼6%) both at 5℃ and 30℃ (Fig. S5b).

**Figure 3.**
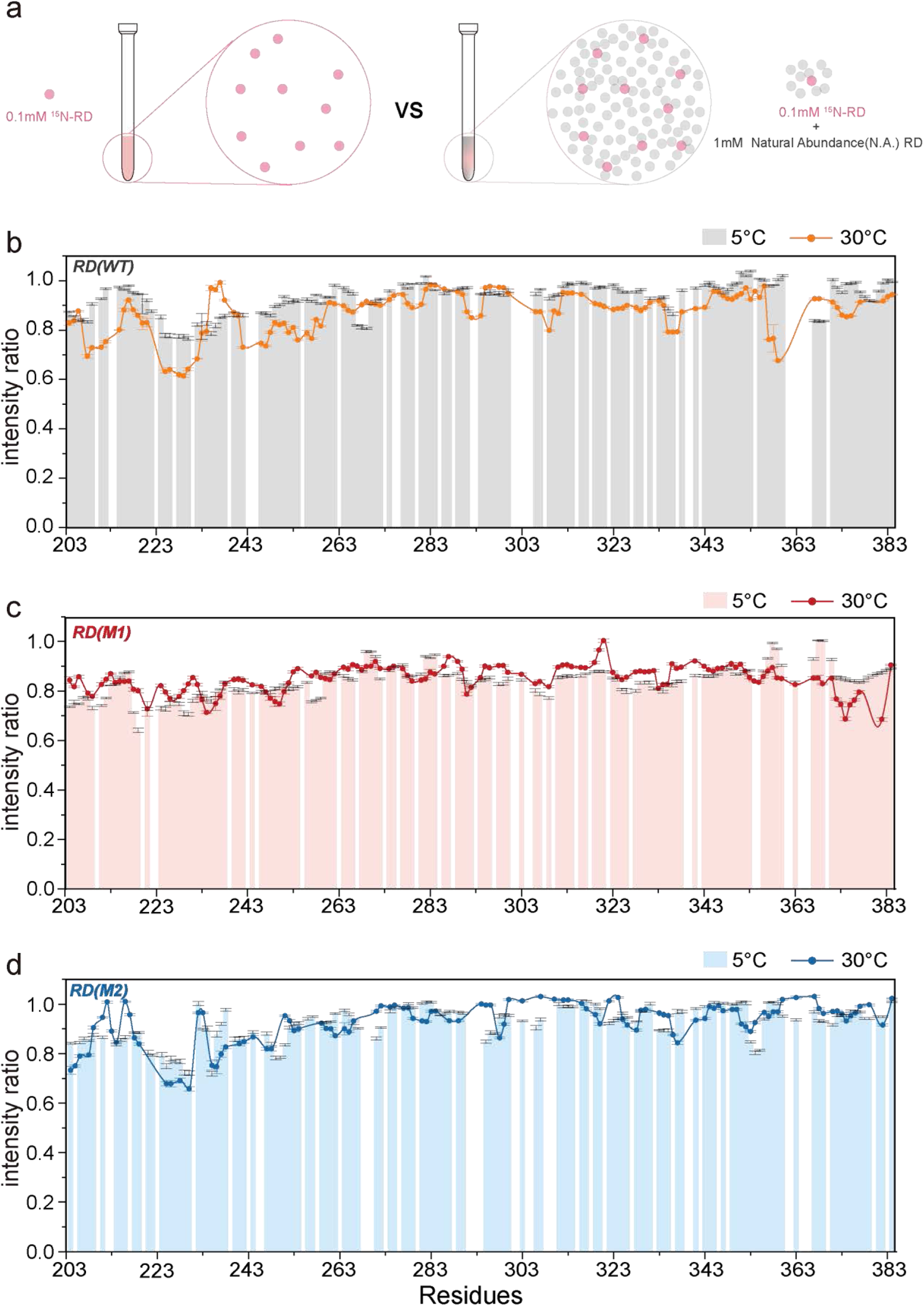
The intermolecular interactions among the RD domains in solution were modulated by temperature and PTMs, as revealed by NMR. **a** Schematic diagram of NMR titration experiments. **b-d** The NMR signal intensity ratio of every residue of the WT RD, M1 and M2 when it was alone or in the presence of 10-fold excess unlabeled counter partner at 5°C and 30°C . The columns represent the intensity ratio at 5°C, while the lines depict the intensity ratio at 30°C , respectively. Points were connected for eye guidance only.

Three pairs of samples are prepared and measured at 5 ℃, 15℃, 25 ℃, and 30℃ for M1, M2, and WT RD. In all cases, we did not observe the chemical shift perturbation but only the changes in the signal intensities. Each residue’s NMR signal intensity ratio was calculated in ^1^H- ^15^N HSQC spectra, revealing interaction sites between RD molecules (Fig. 3 and S6).

At 5 ℃, RD(WT) had a single interaction site located around residues 220-260, while at 15℃, 25℃ and 30℃, more interaction sites emerged and intensities attenuation enhanced, including residues 203-260, 290-310, 313-343 and 355-359 (Fig. 3b and S6a). The spectra of M212 and A329 in WT RD were shown (Fig. S6d and S6e), which were in the abovementioned important interaction regions. The same NMR titration experiments were carried out for M1 and M2 mutants at 5 ℃ to 30 ℃. In contrast to RD(WT), the interaction patterns in M1 and M2 showed no significant change with temperature. Compared to RD(WT), M1 had lower intensity ratios, and more residues in M1 were involved in the interactions (Fig. 3c and S6b), whereas M2 had fewer interaction sites and less signal attenuation (Fig. 3d and S6c). These results indicate that the intermolecular interactions are enhanced in the M1 mutant and weakened in M2. Furthermore, the interaction patterns are consistent with the different phase separation abilities of the three protein versions and support the notion that only RD (WT) undergoes LCST phase separation in this temperature range.

### NMR data-driven coarse grain simulation recaptured the phase-transition behaviors

After revealing the physical chemistry basis of HSF1 phase separation through the mean-field model fitting of the phase diagram, we further delved into the high-resolution mechanism of the HSF1 phase transition and the opposing phase behaviors of M1 and M2. We initially attempted sequence analysis using localCIDER to explore if charge distribution or other sequence parameters would give an intuitive interpretation^50,51^. Unfortunately. our findings indicate that these parameters alone do not fully account for the observed phase behavior (Table S2). Residue-level CG models are particularly useful in uncovering mechanistic details such as critical interactions and crucial sites responsible for driving phase separation in HSF1^52,53^. However, the existing models, such as the hydrophobic scale model with amino acid type-specified force-field parameters, often face limitations in capturing the atomistic details and sequence-encoded biased conformational ensembles, especially for disordered proteins^54,55^. Our attempts to use a few state-of-the-art amino acid type-specific force field parameters^30,31,53,56^ again failed to distinguish the phase behaviors of M1 and M2(Fig. S7). All these suggested that the phase behavior of HSF1 and its mutants is intricately determined by synergistic effects of multiple factors, necessitating a new model that incorporates the collective impact of multi-body interactions among surrounding residues.

In addressing this challenge, we turned to the NMR titration results obtained in Fig. 3. Since the interactions among RD domains were in the slow-exchange regime, the intensity ratios in Fig 3 measured the probability of this residue ‘remaining’ in a ‘free-state’ in the high concentration, thus providing a piece of perfect residue-level structure information similar to those used in data-driven structure protein simulations^57^. Hence, we can compute the simulated NMR intensity ratios based on the simulation trajectory and compare them to the experimental values (See method Section for details). Using the NMR data, we were able to iteratively fine-tune the interaction parameters at each residue of the RD region to satisfy the signal ratios at this site (Fig. 4a and S8), which integrates the entropy effect of solvation and thus bridging the gap between the macroscopic behavior and the molecular-level details. A total of 8 iterations were performed to determine the final set of data-driven force fields. As the iterations progressed, the error declined rapidly, starting at 0.11 and reaching 0.02 by the end (Fig. S8). All three protein sequences were parameterized at 5 ℃, 15 ℃, 25 ℃, and 30 ℃ using the same method.

**Figure 4.**
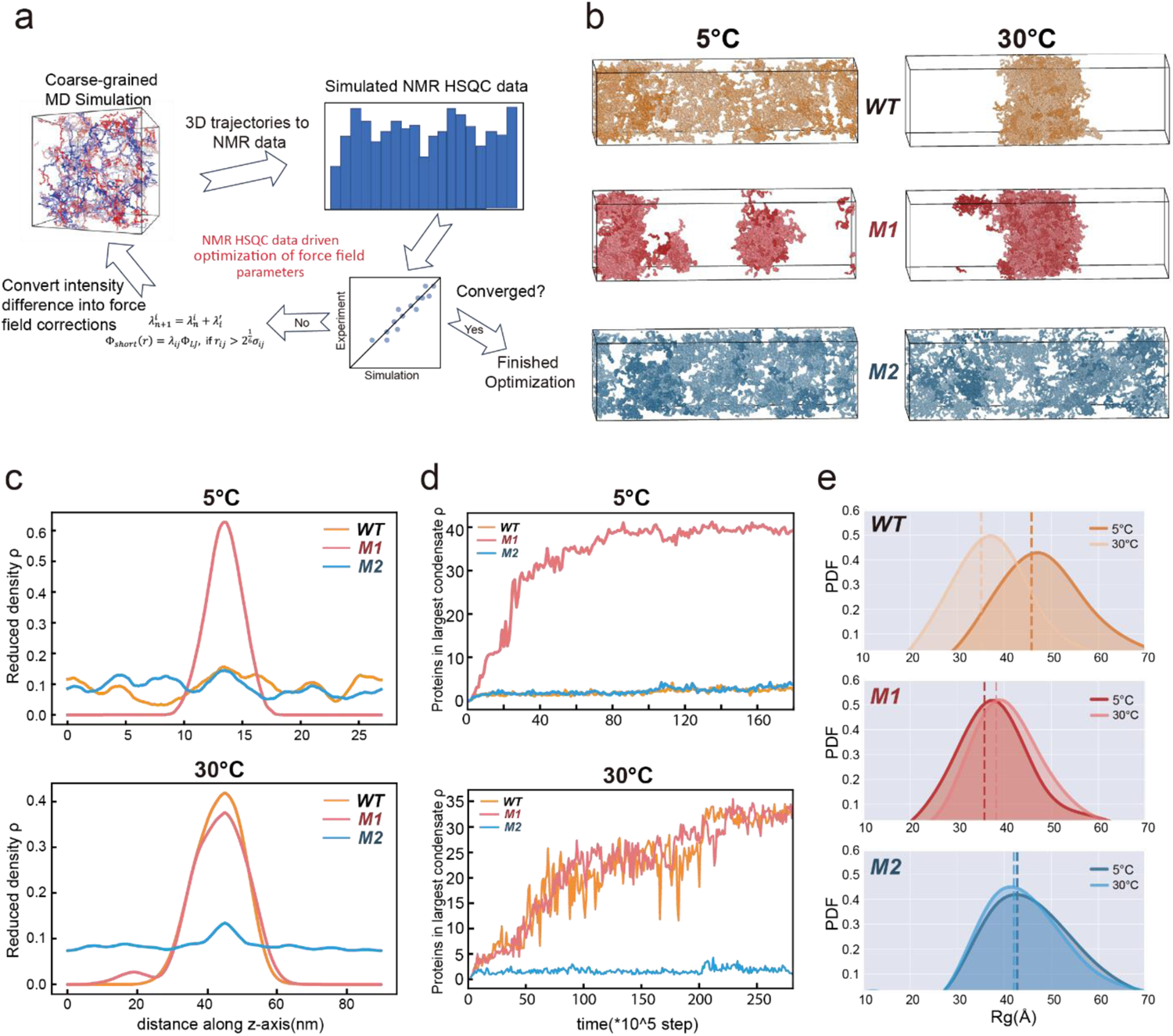
NMR data-driven CG simulation reproduced HSF1 temperature-dependent phase separation and PTM modulations. **a** Illustration of the iterative algorithm for coarse-grained force field optimization. **b** Snapshots of the CG simulation, which reproduced the experimental phase separation observation of LZ1-3-RD at 5°C and 30°C . LZ1-3-RD(M1) and LZ1-3-RD(WT) formed a cluster at 30°C , and LZ1-3-RD(M2) maintained a dispersive distribution. **c** Distributions of the reduced density ρ of WT, M1, and M2 in the CG simulation at 5°C and 30°C . **d** Changes in the protein amounts in the largest condensate ρ of WT, M1, and M2 during CG simulation at 5°C and 30°C . **e** The distribution of the RD domain ensemble’s radius of gyration (Rg) from the CG simulation. The dashed lines indicate the average ensemble values. PDF, Probability Distribution Function.

Based on the NMR data-derived site-specific interaction parameters, the simulations of LZ1-3-RD were successful in replicating experimental observations. Thermodynamic simulations showed that LZ1-3-RD(M1) formed a coexistence phase at all four temperatures (Fig. 4b and c, Fig S9). In contrast, LZ1-3-RD(M2) maintained a dispersive distribution at all four temperatures, consistent with the experimental results (Fig. 1). On the other hand, LZ1-3-RD(WT) formed large condensates at 25 ℃ and 30 ℃ and did not undergo phase transition at 5 ℃ (Fig. 4b and c, Fig. S9). When the temperature was at the intermediate temperature of 15 ℃, the trajectories of wt and M2 did not show obvious differences in the simulation time, while one still can see that wt has a stronger tendency to form instantaneous large clusters than M2 (Fig.S9). Dynamic simulations also indicated that only LZ1-3-RD(M1) formed clusters at 5 ℃, while both LZ1-3-RD(M1) and LZ1-3-RD(WT) formed clusters at 25 ℃ and 30 ℃. Moreover, LZ1-3-RD(M1) had a faster cluster growth than the wild type, as demonstrated in Fig. 4d and Fig. S9. The data-driven model also captured the solely temperature-dependent compactness of wild-type RD with Rg values (Fig. 4e). The data-driven coarse-grained model largely reproduced the experimental observations.

### Specific regions of intermolecular interactions mediate the phase separations of HSF1

Based on the CG simulations, the contact probabilities of each residue participating in intermolecular associations between HSF1 molecules were identified (Fig. 5a). Although no well-defined interaction sites were observed, several stretchers, especially the N-terminal parts, were more frequently involved in interactions in both wild type and M1. Remarkably, the regions with mutations in M1, such as residues 320-343 with 3 mutations in M1, were found to be more involved in the interactions compared with wild-type proteins. Conversely, the N-terminus and the 320-343 region of M2 were less involved in the interactions compared to the wild-type and M1 mutants. These findings suggested that the mutations in M1 and M2 have distinct effects on the intermolecular associations between HSF1 molecules.

**Figure 5.**
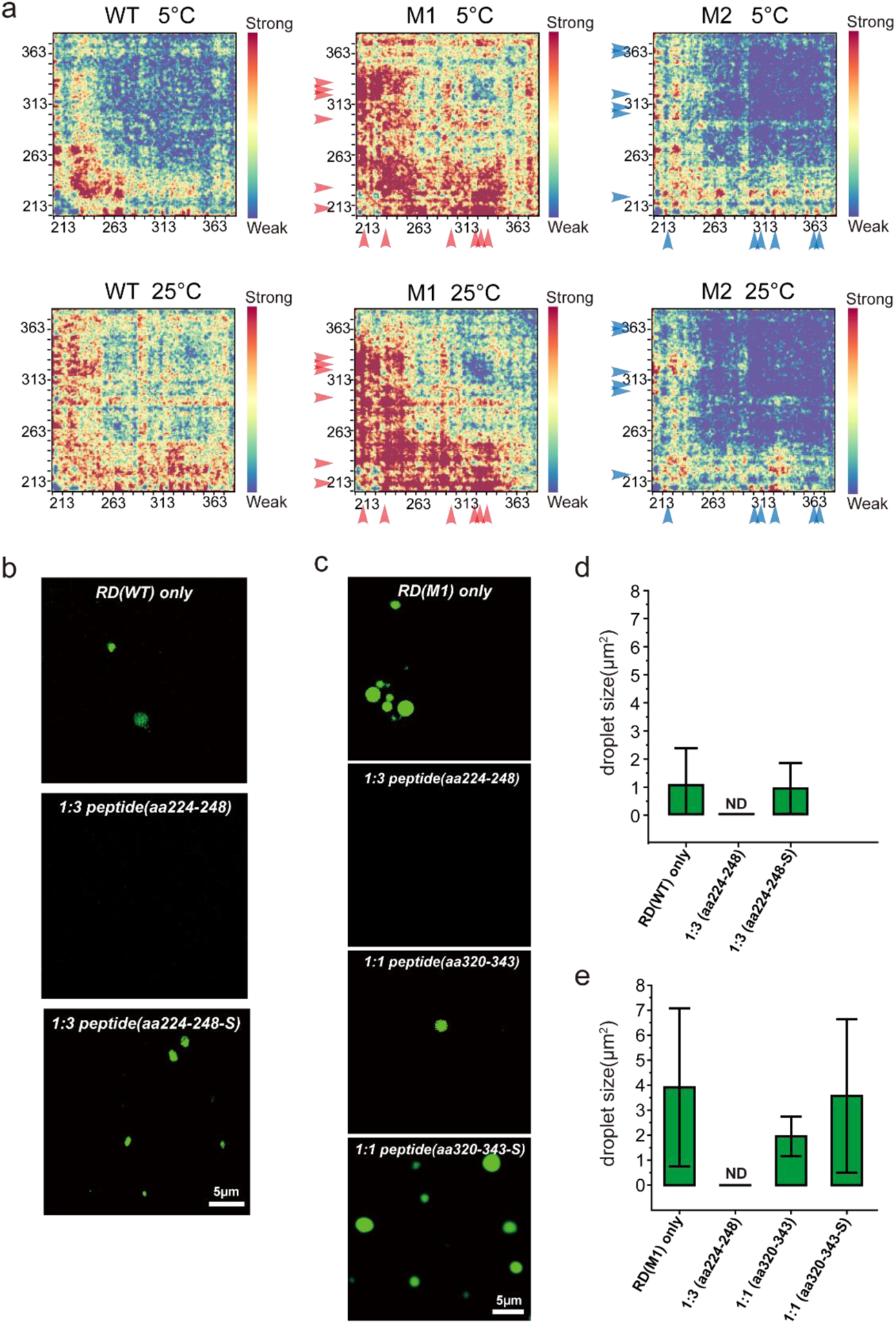
Regions of intermolecular interactions mediate the phase separations of HSF1. **a** Contact maps showing intermolecular interactions between HSF1 molecules from different CG simulation ensembles at 5℃ and 25℃. The mutation sites in M1 and M2 are labeled by red and blue arrows, respectively. **b, c** Peptides originating from the interaction hot spots on contact maps suppress the phase separation of both WT RD and M1 mutants. Scale bars, 5 µm. **d, e.** Droplet sizes from the different treatment groups in **b** and **c**. ND, not determined.

To further validate the crucial role of the intermolecular interactions in phase separation, a “peptide inhibition assay” was carried out. It is rational to assume that excess peptides from the interacting regions would compete with their protein counterparts, reducing the interaction between protein molecules and thus abolishing the phase transition. Two peptides were selected from the regions extensively involved in the intermolecular contacts, as indicated by the contact maps from the CG simulation (Fig. 5a). The peptide derived from the abovementioned N-terminal region (residues 224-248) effectively suppressed the phase separation of both the WT RD (Fig. 5b and 5d) and M1 (Fig. 5c and 5e) mutant at a protein-to-peptide molar ratio of 1:3 at 25 ℃. The peptide originating from the mutation-rich region, residues 320-343, suppressed phase separation more efficiently, significantly reducing droplets even at a protein: peptide ratio of 1:1 (Fig. 5c and 5e). Two peptides were designed as negative controls to check the specificity of interactions, whose hydrophobic residues were mutated to Ser but with the same net charge and length. These two peptides, dubbed as aa224-248-S and aa320-343-S(Fig. S10a and S10b), failed to inhibit the phase transition of wt and M1 RD domains (Fig. 5b, 5c, 5d and 5e). The inhibitory effects of these two peptides confirmed the significance of residues 224-248 and residue 320-343 regions in the phase separation of HSF1. These results further validate the accuracy of CG simulations.

### The interaction regions govern the phase-transition temperature of HSF1, which equips the physiology of animal body temperature

The phase transition temperatures of the HSF1 from several animal species are found to be different. We compared the sequences for HSF1 from the abovementioned animal species to find the key region that determines the phase transition temperatures. Sequence alignment revealed that the LZ1-3-RD-LZ4 regions were largely conserved among these species, with the primary deviations between residue 320 and 360 (Fig. 6a). This region of human HSF1 is critical for phase separation and is more involved in intermolecular interactions when temperature rises, as demonstrated in peptide inhibition assay and contact maps from CG simulation (Fig. 5). The zebrafish HSF1 shows additional differences in the LZ4 region, which may affect the interaction between leucine zippers.

**Figure 6.**
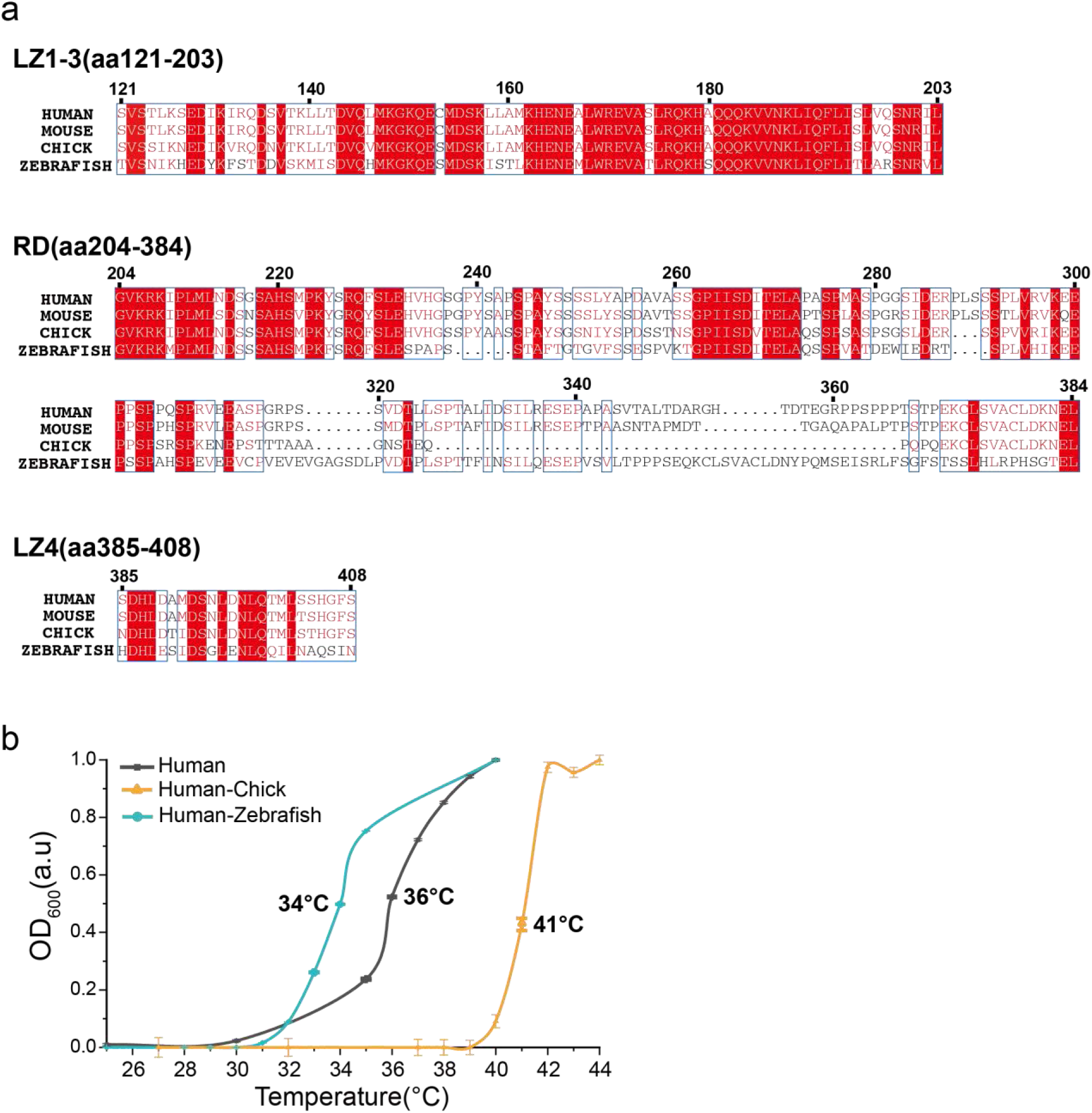
The regions with temperature-dependent intermolecular interactions differ among the animals with different body temperatures and determine the phase transition temperatures. **a.** The sequence alignment of the LZ1-3-RD-LZ4 regions from HSF1 of humans, mice, chicken, and zebrafish showed the main species-specific differences between residue 320 and 360, matching with the regions with a temperature-dependent interaction pattern in Figure 6a. **b** The turbidity-temperature curves of human HSF1(LZ1-3-RD-LZ4) with the 320-360 regions replaced by the counterparts from chicken and zebrafish were compared to the wild-type protein. Swapping the region corresponding to human residues 320-360 changed the LLSP temperatures, providing further evidence of the importance of these specific regions in determining phase transition temperatures.

To validate the key roles of the residue 320-360 regions in determining the HSF1 phase separation temperature, we carried out two “sequence-swapping” assays. Consistent with our prediction, the replacement of the region of residues 320-360 of human HSF1 by the corresponding region from zebrafish, the critical phase separation temperature for the “human-zebrafish HSF1” hybrid got lower (Fig. 6b). On the other hand, if this region was replaced by its counterparts from the chicken HSF1, the “human-chicken HSF1” hybrid’s phase separation initiated at a higher temperature than the wild-type protein (Fig. 6b). These results demonstrated that the interaction sites at residues 320-360 determine the transition temperatures.

## Discussion

Current research has demonstrated that HSF1 undergoes an LCST-type phase separation, and its phase separation temperature matches the body temperature of animals. This discovery has led to the proposal that HSF1 functions as an “in-cell thermometer” by detecting temperature changes and reacting to heat shock through phase separation and the critical amino acids underlying HSF1 phase separation constitute the temperature code. While there have been reports on protein LCST phase transitions before^58,59^, the temperature-dependent phase separation property of HSF1 carries significant physiological implications crucial for its role in HSR^13,17,19,60^. Our study, which integrated biochemistry, biophysics, and simulation methods, also successfully dissected the mechanism of phase transition and PTM modulations down to the residue level and has experimentally demonstrated that the critical interaction regions can determine the phase separation temperature.

The LCST-type phase separation of HSF1 we found here provides a new molecular mechanism of HSF1 activation in heat shock. The previously proposed HSF1 activation mechanisms, such as leucine zippers oligomerization ^11^, only modulate the temperature-dependent phase separation rather than the root cause. Upon initiating the heat shock response, the LZ4 domain dissociates from LZ1-3, enabling LZ1-3 to promote intermolecular oligomerization, which triggers phase separation by elevating the local protein concentration and the multivalency of interaction (Fig. 7). Without the LZ4 motif, phase separation will occur at a much lower temperature, such as 25 ℃ (Fig. 1d and 1e). The collaboration of the phase separation nature from the RD domain and oligomerization properties of the LZ motifs restricts phase separation to temperatures around or above physiological levels, with a marked transition once this threshold is reached. These abrupt changes in phase transition behavior render HSF1 a highly sensitive indicator of heat shock.

**Figure 7.**
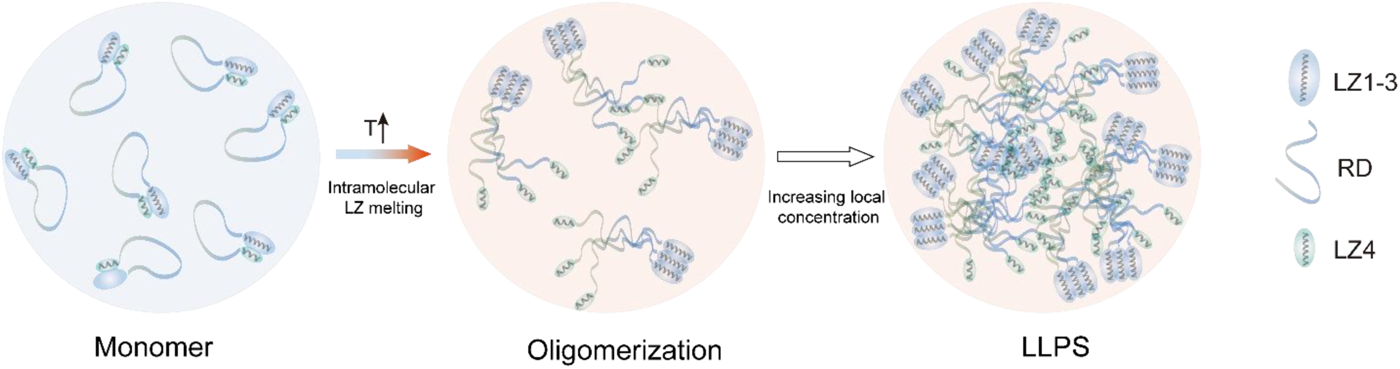
Schematic diagram of the HSF1 activation mechanism, which involves the synergistic effects of phase separation mediated by the RD domain and oligomerization by LZ1-3. At low temperatures, the LZ1-3 binds with LZ-4, which keeps the HSF1 as a monomer, as shown in the left panel. As the temperature increases, the LZ4 dissociates from LZ1-3, enabling HSF1 oligomerizing via LZ1-3, as shown in the middle panel. Oligomerization increases the local concentration, thus initiating the phase separation.

The finding of this study established a correlation between the phase separation temperatures of the HSF1 from different animal species and their respective body temperatures, highlighting the physiological relevance of the HSF1 phase separation. Notably, we observed that sequence differences among species are predominantly located in the region where interactions change with temperature, which is a key determinant of phase separation temperatures. These findings further confirm the accuracy of the CG simulation ensembles in studying the mechanisms underlying HSF1 phase separation.

The interaction sites identified in this study align with those reported in previous functional studies. The N-terminal region of the RD domain has previously been shown to be recognized by the chaperone, which helps to disperse the aggregates of HSF1^61^. Our results also indicate that the residue 320-343 region of the M1 RD domain, which includes three mutations (S320D, S326D, and S333D), is emerging as a new interaction hub. It was reported that phosphorylation of these three sites is correlated with the upregulation of HSF1 activity ^62–64^.

The findings of our study also revealed that PTM is not a prerequisite for phase separation at heat shock temperatures but instead serves as a modulation mechanism. Previous studies have suggested that phosphorylation is necessary for enabling phase separation in HSF1^13^. However, such studies were conducted at room temperature and did not mimic the physiological heat shock temperatures required for the phase separation of the wild-type proteins.

The PTMs mimicked by M1 and M2 are phosphorylation and acetylation, which not only introduce negative charges but also reduce the hydrophobicity around that site. The mean field model showed that the entropy changes associated with surface-restricted water are crucial factors in the LCST-type behavior, which varied upon phosphorylation due to the difference of the hydrophobic surfaces. The (mimicked) phosphorylation reduced the hydrophobicity, making the phase separation less temperature-dependent in the cases of M1 and M2. When we compensate for the hydrophobicity loss due to phosphorylation by introducing more hydrophobic residues around the phosphorylation sites, the LCST-type behavior also appeared in the M1 mutants. While it is known in physical chemistry that the entropically driven release of solvent is the origin of LCST phase separation^47^, explicitly underlying its importance to sequence determinants of biological condensates was rarely discussed despite a few works^59,65,66^.

The potential effects of PTMs on intermolecular interactions were analyzed at the residue level. The changes in interaction patterns cannot be explained solely by the charges of involved fragments, indicating the presence of other factors, such as hydrophobicity. While previous research has reported the effects of phosphorylation-induced hydrophobic interactions on long-range intramolecular interactions within the intrinsically disordered domains^67^, our study here focused on inter-molecular interactions at physiological heat shock temperatures. Nonetheless, a more detailed understanding of the influences at the atomic level will require further investigation.

In the current study, extensive interaction information was extracted from NMR data and used to optimize a residue-based force field. Our novel approach significantly improved the accuracy of the force field. While the knowledge-based model has been widely used for folded proteins with X-ray scattering data^65,68,69^, our work provides a complementary algorithm for studying interactions in disordered proteins. Furthermore, the CG simulation also independently reproduced the changes in chain compactness, which is correlated to the phase separation propensity of IDRs^42,43^. It is important to note that without knowledge of the interaction sites and strength from NMR titration, the simulation using existing force fields failed to reproduce the phase separation, which was mainly due to the inability of the available force field to accurately represent the slight difference caused by a few mutations in the sequences (Fig. S7).

In sum, this study presents a comprehensive analysis of the phase separation mechanism of HSF1 and identifies a species-specific temperature code of HSF1 accounting for the physiology of individual species, which offers a novel insight into the temperature sensing of phase separation. The temperature-dependence phase separation behaviors of HSF1 make it a promising tool for probing the temperature-dependent phase separation driven events in cell biology. The great excitement ahead is to delineate the molecular-level changes in the dynamic properties of HSF1 droplets and the details of how condensed other proteins disperse HSF1 foci during heat shock responses in living cells.

## Acknowledgment

We thank the staff, Dr. Guangfeng Liu, from BL19U2 beamline of the National Facility for Protein Science in Shanghai (NFPS) at Shanghai Synchrotron Radiation Facility, for assistance during SAXS data collection. This work was supported by grants from the National Key R&D Program (2019YFA0508403) and the National Natural Science Foundation of China (31971128, 32090044), the Strategic Priority Research Program of the Chinese Academy of Sciences Grant No. XDB 37040202, CAS Project for Young Scientists in Basic Research Grant No. YSBR-068.

## References

1. Anckar, J. & Sistonen, L. Regulation of HSF1 function in the heat stress response: implications in aging and disease. Annu Rev Biochem 80, 1089–115 (2011).

2. Zheng, X. et al. Dynamic control of Hsf1 during heat shock by a chaperone switch and phosphorylation. Elife 5(2016).

3. Kijima, T. et al. HSP90 inhibitors disrupt a transient HSP90-HSF1 interaction and identify a noncanonical model of HSP90-mediated HSF1 regulation. Sci Rep 8, 6976 (2018).

4. Li, J., Labbadia, J. & Morimoto, R.I. Rethinking HSF1 in Stress, Development, and Organismal Health. Trends Cell Biol 27, 895–905 (2017).

5. Mendillo, M.L. et al. HSF1 drives a transcriptional program distinct from heat shock to support highly malignant human cancers. Cell 150, 549–62 (2012).

6. Meng, L., Gabai, V.L. & Sherman, M.Y. Heat-shock transcription factor HSF1 has a critical role in human epidermal growth factor receptor-2-induced cellular transformation and tumorigenesis. Oncogene 29, 5204–13 (2010).

7. Dai, C., Whitesell, L., Rogers, A.B. & Lindquist, S. Heat shock factor 1 is a powerful multifaceted modifier of carcinogenesis. Cell 130, 1005–18 (2007).

8. Goetzl, E.J. et al. Low neural exosomal levels of cellular survival factors in Alzheimer’s disease. Ann Clin Transl Neurol 2, 769–73 (2015).

9. Gomez-Pastor, R. et al. Abnormal degradation of the neuronal stress-protective transcription factor HSF1 in Huntington’s disease. Nat Commun 8, 14405 (2017).

10. Gomez-Pastor, R., Burchfiel, E.T. & Thiele, D.J. Regulation of heat shock transcription factors and their roles in physiology and disease. Nat Rev Mol Cell Biol 19, 4–19 (2018).

11. Hentze, N., Le Breton, L., Wiesner, J., Kempf, G. & Mayer, M.P. Molecular mechanism of thermosensory function of human heat shock transcription factor Hsf1. Elife 5(2016).

12. Xu, Y.M., Huang, D.Y., Chiu, J.F. & Lau, A.T. Post-translational modification of human heat shock factors and their functions: a recent update by proteomic approach. J Proteome Res 11, 2625–34 (2012).

13. Zhang, H. et al. Reversible phase separation of HSF1 is required for an acute transcriptional response during heat shock. Nat Cell Biol 24, 340–352 (2022).

14. Lafontaine, D.L.J., Riback, J.A., Bascetin, R. & Brangwynne, C.P. The nucleolus as a multiphase liquid condensate. Nat Rev Mol Cell Biol 22, 165–182 (2021).

15. Gibson, B.A. et al. Organization of Chromatin by Intrinsic and Regulated Phase Separation. Cell 179, 470–484 e21 (2019).

16. Chong, S. et al. Tuning levels of low-complexity domain interactions to modulate endogenous oncogenic transcription. Mol Cell 82, 2084–2097 e5 (2022).

17. Cotto, J., Fox, S. & Morimoto, R. HSF1 granules: a novel stress-induced nuclear compartment of human cells. J Cell Sci 110 ( Pt 23), 2925–34 (1997).

18. Jolly, C., Morimoto, R., Robert-Nicoud, M. & Vourc’h, C. HSF1 transcription factor concentrates in nuclear foci during heat shock: relationship with transcription sites. J Cell Sci 110 (Pt 23), 2935–41 (1997).

19. Gaglia, G. et al. HSF1 phase transition mediates stress adaptation and cell fate decisions. Nat Cell Biol 22, 151–158 (2020).

20. Martin, E.W. et al. Valence and patterning of aromatic residues determine the phase behavior of prion-like domains. Science 367, 694–699 (2020).

21. Kim, T.H. et al. Phospho-dependent phase separation of FMRP and CAPRIN1 recapitulates regulation of translation and deadenylation. Science 365, 825–829 (2019).

22. Ibáñez de Opakua, A., et al. Molecular interactions of FG nucleoporin repeats at high resolution. Nat Chem 14, 1278–1285 (2022).

23. Paloni, M., Bailly, R., Ciandrini, L. & Barducci, A. Unraveling Molecular Interactions in Liquid–Liquid Phase Separation of Disordered Proteins by Atomistic Simulations. The Journal of Physical Chemistry B 124, 9009–9016 (2020).

24. Collepardo-Guevara, R. et al. Chromatin Unfolding by Epigenetic Modifications Explained by Dramatic Impairment of Internucleosome Interactions: A Multiscale Computational Study. Journal of the American Chemical Society 137, 10205–10215 (2015).

25. Guilln-Boixet, J. et al. RNA-Induced Conformational Switching and Clustering of G3BP Drive Stress Granule Assembly by Condensation. Cell 181, 346–361.e17 (2020).

26. McCarty, J., Delaney, K.T., Danielsen, S.P.O., Fredrickson, G.H. & Shea, J.E. Complete Phase Diagram for Liquid-Liquid Phase Separation of Intrinsically Disordered Proteins. Journal of Physical Chemistry Letters 10, 1644--1652 (2019).

27. Lin, Y.H. & Chan, H.S. Phase Separation and Single-Chain Compactness of Charged Disordered Proteins Are Strongly Correlated. Biophysical Journal 112, 2043--2046 (2017).

28. Das, S., Lin, Y.-H., Vernon, R.M., Forman-Kay, J.D. & Chan, H.S. Comparative roles of charge, π, and hydrophobic interactions in sequence-dependent phase separation of intrinsically disordered proteins. Proceedings of the National Academy of Sciences 117, 28795–28805 (2020).

29. Li, L.G. & Hou, Z. Theoretical modelling of liquid-liquid phase separation: from particle-based to field-based simulation. Biophys Rep 8, 55–67 (2022).

30. Regy, R.M., Thompson, J., Kim, Y.C. & Mittal, J. Improved coarse-grained model for studying sequence dependent phase separation of disordered proteins. Protein Sci 30, 1371–1379 (2021).

31. Tesei, G., Schulze, T.K., Crehuet, R. & Lindorff-Larsen, K. Accurate model of liquid–liquid phase behavior of intrinsically disordered proteins from optimization of single-chain properties. Proceedings of the National Academy of Sciences 118, e2111696118 (2021).

32. Tejedor, A.R. et al. Protein structural transitions critically transform the network connectivity and viscoelasticity of RNA-binding protein condensates but RNA can prevent it. Nature Communications 13, 5717 (2022).

33. Alexander, E.C. et al. TDP-43 α-helical structure tunes liquid–liquid phase separation and function. Proceedings of the National Academy of Sciences 117, 5883–5894 (2020).

34. Bremer, A. et al. Deciphering how naturally occurring sequence features impact the phase behaviours of disordered prion-like domains. Nat Chem 14, 196–207 (2022).

35. Najbauer, E.E., Ng, S.C., Griesinger, C., Gorlich, D. & Andreas, L.B. Atomic resolution dynamics of cohesive interactions in phase-separated Nup98 FG domains. Nat Commun 13, 1494 (2022).

36. Babu, M., Favretto, F., Rankovic, M. & Zweckstetter, M. Peptidyl Prolyl Isomerase A Modulates the Liquid-Liquid Phase Separation of Proline-Rich IDPs. J Am Chem Soc 144, 16157–16163 (2022).

37. Murthy, A.C. et al. Molecular interactions underlying liquid−liquid phase separation of the FUS low-complexity domain. Nature Structural & Molecular Biology 26, 637–648 (2019).

38. Gomes, G.-N.W. et al. Conformational Ensembles of an Intrinsically Disordered Protein Consistent with NMR, SAXS, and Single-Molecule FRET. Journal of the American Chemical Society 142, 15697–15710 (2020).

39. Refinetti, R. The circadian rhythm of body temperature. Front Biosci (Landmark Ed) 15, 564–94 (2010).

40. Piestun, Y., Druyan, S., Brake, J. & Yahav, S. Thermal manipulations during broiler incubation alter performance of broilers to 70 days of age. Poult Sci 92, 1155–63 (2013).

41. López-Olmeda, J.F. & Sánchez-Vázquez, F.J. Thermal biology of zebrafish (Danio rerio). Journal of Thermal Biology 36, 91–104 (2011).

42. Lin, Y.-H. & Chan, H.S. Phase Separation and Single-Chain Compactness of Charged Disordered Proteins Are Strongly Correlated. Biophysical Journal 112, 2043–2046 (2017).

43. Li, L. & Hou, Z. Crosslink-Induced Conformation Change of Intrinsically Disordered Proteins Have a Nontrivial Effect on Phase Separation Dynamics and Thermodynamics. The Journal of Physical Chemistry B 127, 5018–5026 (2023).

44. Quiroz, F.G. & Chilkoti, A. Sequence heuristics to encode phase behaviour in intrinsically disordered protein polymers. Nature Materials 14, 1164--1171 (2015).

45. Quiroz, F.G. et al. Intrinsically disordered proteins access a range of hysteretic phase separation behaviors. Science Advances 5, 1--12 (2019).

46. 46. Kinetics of liquid-liquid phase separation in protein solutions exhibiting LCST phase behavior studied by time-resolved USAXS and VSANS. Soft Matter 12, 9334–9341 (2016).

47. Matsuyama, A. & Tanaka, F. Theory of solvation-induced reentrant phase separation in polymer solutions. Physical review letters 65, 341 (1990).

48. Türkaydin, B. et al. From head to tail - Atomistic mechanism of long-range coupling from the cytosolic sensor domain to the selectivity filter in TREK K 2P channels. bioRxiv, 2023.10.06.561191 (2023).

49. Winter, H., Huber, J.L. & Huber, S.C. Membrane association of sucrose synthase: changes during the graviresponse and possible control by protein phosphorylation. FEBS Lett 420, 151–5 (1997).

50. Martin, E.W. et al. Sequence Determinants of the Conformational Properties of an Intrinsically Disordered Protein Prior to and upon Multisite Phosphorylation. Journal of the American Chemical Society 138, 15323–15335 (2016).

51. Das, R.K. & Pappu, R.V. Conformations of intrinsically disordered proteins are influenced by linear sequence distributions of oppositely charged residues. Proceedings of the National Academy of Sciences 110, 13392–13397 (2013).

52. Ma, Y. et al. Nucleobase Clustering Contributes to the Formation and Hollowing of Repeat-Expansion RNA Condensate. J Am Chem Soc 144, 4716–4720 (2022).

53. 53. Mammen Regy, R., Zheng, W. & Mittal, J. Chapter One - Using a sequence-specific coarse-grained model for studying protein liquid–liquid phase separation. in Methods in Enzymology, Vol. 646 (ed. Keating, C.D.) 1–17 (Academic Press, 2021).

54. Shrestha, U.R. et al. Generation of the configurational ensemble of an intrinsically disordered protein from unbiased molecular dynamics simulation. Proceedings of the National Academy of Sciences 116, 20446–20452 (2019).

55. Kulkarni, P. et al. Intrinsically disordered proteins: Ensembles at the limits of Anfinsen’s dogma. Biophysics Reviews 3(2022).

56. Dannenhoffer-Lafage, T. & Best, R.B. A Data-Driven Hydrophobicity Scale for Predicting Liquid–Liquid Phase Separation of Proteins. The Journal of Physical Chemistry B 125, 4046–4056 (2021).

57. Go, N. Theoretical studies of protein folding. Annual review of biophysics and bioengineering 12, 183–210 (1983).

58. Riback, J.A. et al. Stress-Triggered Phase Separation Is an Adaptive, Evolutionarily Tuned Response. Cell 168, 1028–1040 e19 (2017).

59. Chen, S. & Wang, Z.G. Using Implicit-Solvent Potentials to Extract Water Contributions to Enthalpy-Entropy Compensation in Biomolecular Associations. J Phys Chem B 127, 6825–6832 (2023).

60. Chowdhary, S., Kainth, A.S., Paracha, S., Gross, D.S. & Pincus, D. Inducible transcriptional condensates drive 3D genome reorganization in the heat shock response. Mol Cell 82, 4386–4399 e7 (2022).

61. Kmiecik, S.W., Le Breton, L. & Mayer, M.P. Feedback regulation of heat shock factor 1 (Hsf1) activity by Hsp70-mediated trimer unzipping and dissociation from DNA. The EMBO Journal 39, e104096 (2020).

62. Guettouche, T., Boellmann, F., Lane, W.S. & Voellmy, R. Analysis of phosphorylation of human heat shock factor 1 in cells experiencing a stress. BMC Biochem 6, 4 (2005).

63. Murshid, A. et al. Protein kinase A binds and activates heat shock factor 1. PLoS One 5, e13830 (2010).

64. Sourbier, C. et al. Englerin A stimulates PKCθ to inhibit insulin signaling and to simultaneously activate HSF1: pharmacologically induced synthetic lethality. Cancer Cell 23, 228–37 (2013).

65. Da Vela, S. et al. Kinetics of liquid-liquid phase separation in protein solutions exhibiting LCST phase behavior studied by time-resolved USAXS and VSANS. Soft Matter 12, 9334–9341 (2016).

66. Matsarskaia, O. et al. Tuning phase transitions of aqueous protein solutions by multivalent cations. Physical Chemistry Chemical Physics 20, 27214–27225 (2018).

67. Du, Z. et al. Phosphorylation modulates estrogen receptor disorder by altering long-range hydrophobic interactions. bioRxiv, 2023.07.14.548966 (2023).

68. Kolakofsky, D., Kowalinski, E. & Cusack, S. A structure-based model of RIG-I activation. RNA 18, 2118–2127 (2012).

69. Lane, T.J., Shukla, D., Beauchamp, K.A. & Pande, V.S. To milliseconds and beyond: challenges in the simulation of protein folding. Current Opinion in Structural Biology 23, 58–65 (2013).

70. Thompson, J.D., Higgins, D.G. & Gibson, T.J. CLUSTAL W: improving the sensitivity of progressive multiple sequence alignment through sequence weighting, position-specific gap penalties and weight matrix choice. Nucleic Acids Res 22, 4673–80 (1994).

71. Robert, X. & Gouet, P. Deciphering key features in protein structures with the new ENDscript server. Nucleic Acids Research 42, W320–W324 (2014).

72. Hopkins, J.B., Gillilan, R.E. & Skou, S. BioXTAS RAW: improvements to a free open-source program for small-angle X-ray scattering data reduction and analysis. J Appl Crystallogr 50, 1545–1553 (2017).

73. Delaglio, F. et al. NMRPipe: a multidimensional spectral processing system based on UNIX pipes. J Biomol NMR 6, 277–93 (1995).

74. Lee, W., Tonelli, M. & Markley, J.L. NMRFAM-SPARKY: enhanced software for biomolecular NMR spectroscopy. Bioinformatics 31, 1325–7 (2015).

75. Kjaergaard, M. & Poulsen, F.M. Sequence correction of random coil chemical shifts: correlation between neighbor correction factors and changes in the Ramachandran distribution. Journal of Biomolecular NMR 50, 157–165 (2011).

76. Camilloni, C., De Simone, A., Vranken, W.F. & Vendruscolo, M. Determination of Secondary Structure Populations in Disordered States of Proteins Using Nuclear Magnetic Resonance Chemical Shifts. Biochemistry 51, 2224–2231 (2012).

77. Ashbaugh, H.S. & Hatch, H.W. Natively Unfolded Protein Stability as a Coil-to-Globule Transition in Charge/Hydropathy Space. Journal of the American Chemical Society 130, 9536–9542 (2008).

